# Shade avoidance restricts soybean breeding progress and increases herbivore susceptibility

**DOI:** 10.1101/2025.09.24.678302

**Authors:** Lukas Vonmetz, Emanuel B. Kopp, Simon Jäggi, Pascal A. Niklaus, Samuel E. Wuest

## Abstract

High planting densities expose field crops to competition for light, which typically induces shade avoidance responses such as stem elongation. While adaptive in natural environments, these responses can lower yield and increase susceptibility to stress in agricultural systems. We tested whether soybean breeding over the past century has altered shade avoidance and associated trade-offs. Twenty-one Canadian cultivars released between 1922 and 2018 were grown in pots under either control or shade-avoidance-inducing light conditions, achieved by altering reflected light spectrum without reducing photosynthetic radiation. Plants exposed to shade-avoidance inducing light grew taller and suffered greater thrips damage, consistent with expectations of increased stem elongation and reduced defense. More recent cultivars showed higher susceptibility to thrips than older ones. Breeding progress in seed yield was driven largely by greater biomass allocation to seeds and a reduction in branching. However, under shade-inducing light, the yield improvements were smaller, pointing to shade avoidance as a limiting factor. Our results indicate that while soybean breeding has improved yield and shifted morphology towards ideotypes suited for high-density stands, persistent shade avoidance responses constrain breeding progress and increase herbivore susceptibility. Breeding strategies that reduce sensitivity to neighbor cues may therefore improve soybean productivity and resilience.

## Introduction

Densely populated environments impose strong selection on plant traits that improve access to limiting resources, particularly light. Shade avoidance responses thus evolved as adaptive strategy to avoid shade and more efficiently compete for light (Schmitt 1997; Schmitt et al. 2003); this includes stem and petiole elongation, leaf angle adjustment, reduced branching or shifts in phenology (Ballaré et al. 1990; Franklin 2008; Pierik and De Wit 2014; De Wit et al. 2016; Gruntman et al. 2017a). These responses are primarily triggered by a reduction in the ratio of red to far-red (R:FR) light, which signals the presence of neighbouring vegetation (Ballaré et al. 1990). While advantageous for individual fitness in competitive environments, these responses often involve resource allocation trade-offs, and can result in decreased allocation to storage and reproduction or increase susceptibility to lodging (Schmitt et al. 1995; Franklin and Whitelam 2005)

In agricultural systems, plants are typically grown in uniform, high-density stands where overall yield, rather than individual performance, determines success (Duvick et al. 2003). In this context, shade avoidance responses - while adaptive in nature - can reduce yield by reallocating resources from harvestable organs toward elongation or support (Boccalandro et al. 2003; Carriedo et al. 2016). This notion is a core concept of “Darwinian Agriculture” (also known as “Evolutionary Agroecology”), and suggests that traits which evolved to enhance individual competitiveness may conflict with collective crop performance (Denison 2012; Weiner 2019). This view challenges the assumption that the breeding progress leads to an “improvement” of crops and posits that evolutionary trade-off persist in modern breeding programs and may lead to conflicting selective responses, depending on the breeding scheme chosen.

Breeding of field crops typically involves a phase early in the breeding cycle predominantly characterized by individual-level selection (e.g. during mass selection), followed by a phase of predominant selection at the level of groups of highly related individuals (e.g. rows or plots of inbred progenies, leading to multi-level selective regimes; Murphy et al. 2017). This may lead to conflicting selection pressures on “selfish” traits: early individual-level selection favours selfish traits, as competitive individuals stand out against their less competitive neighbours and increase in frequency. Conversely, later group selection favours less competitive traits. Despite this conflict, the analysis of historic series of cultivars released over several decades has revealed that with breeding for higher yields, other traits that are specifically beneficial for high-density groups also increased: vertical leaf angles, steeper roots, higher water and nitrogen use efficiency, etc., (Costa Netto et al. 2025; Duvick et al. 2004; York et al. 2015; Zhu et al. 2019, 2022). At the same time, breeders have also developed idealized plant models, originally denoted as “communal” ideotypes, that express such traits benefiting the group (Donald 1968). A well-known example is the reduction in plant height in wheat (Reitz and Salmon 1968) and rice (Jennings 1964), and recently maize, which increases yield potential under high densities (Kosola et al. 2023). Together, these phenotypic changes therefore reflect deliberate or inadvertent shifts away from competitive traits and towards traits that support collective performance (Denison 2012; Weiner 2019). However, current breeding practises do not guarantee that traits that benefit the group at the expense of individual fitness will automatically be favoured during selection.

From this perspective, limiting shade avoidance as a selfish response to neighbour plants could improve stand-level yield (Golan et al. 2023; Liu et al. 2020; Li et al. 2023). Shade avoidance is a plastic trait, however, and compared to other, often more fixed, traits, relatively less is known about how it has changed with crop breeding progress (Wille et al. 2017). In wheat, genetic changes that reduce plant height indirectly also reduce shade avoidance responses (Colombo et al. 2022; Golan et al. 2024). While there is evidence for genetic variation in shade avoidance response in soybean (Gong et al. 2015), it remains unclear whether the responses of modern soybean cultivars differ from those of historical cultivars.

Studying shade avoidance responses is complicated by the fact that plants can respond to different aspects of the light environment, such as changes in light levels and changes in light spectrum (Ballaré and Pierik 2017). Changes in the light environment of a plant facing a competitor is also typically confounded with various types of resource competition, such as belowground competition for soil nutrients, which also influences plant growth (Murphy and Dudley 2007; Kopp et al. 2025). To manipulate the spectral properties of light, filters are placed above plants, but these typically also attenuate light intensity, making it harder to isolate the specific role of light level and light spectrum.

Here, we studied the effect of breeding in soybean on shade avoidance responses using 21 cultivars released between 1922 to 2018. We positioned light filters beneath plants to manipulate the spectrum of reflected light, specifically the red to far red ratio and amounts of blue light. Because we did not manipulate incident light, levels of radiation available for photosynthesis remained unchanged (Green-Tracewicz et al. 2011). Since we grew single plants in pots, the effect of light quality changes also was not conflated with effects of belowground competition. Specifically, we were interested whether modern soybean cultivars exhibit reduced shade avoidance, which are likely adaptations to higher planting densities and improved group-level yield.

## Materials and Methods

### Shade avoidance experiment

In summer 2024, a pot experiment was performed in Wädenswil (N 47°13’19.7” E 8°40’8.4”) to investigate shade avoidance responses in a set of 21 early-maturing soybean cultivars released in Canada between 1922 to 2018 (Supplementary Materials, Table S1). Seeds were germinated in small pots (0.29 l, filled with FloraDur A-Block, Floragard Vertriebs-GmbH, Germany) in the glasshouse (night: 16°C-18°C, day: 22°C-24°C, photoperiod: 12 h). Pots were arranged with 18 cm distance from each other, arranged on four large tables. The pots were covered by a large plastic sheet with holes allowing plants to grow through. Two tables were covered with a clear plastic sheet not altering the spectrum of the reflected light (control treatment; Clear 130, LEE Filters, UK). The other two tables were covered with a green plastic sheet that reduced the red to far red ratio of reflected light (shade-avoidance inducing treatment, SAI; Fern Green 122, LEE Filters, UK; (Chen et al. 2023; Wille et al. 2017; Gruntman et al. 2017b; Golan et al. 2024). After 10 days, twelve plants per cultivar and treatment were transplanted into 3 liter pots (21 cultivars x 2 treatments x 12 replicates = 504 pots) filled with a special mixture of soybean soil (RICOTER Erdaufbereitung AG, Frauenfeld, Switzerland: 25% garden soil max. 15 mm; 60% white peat 0-30 mm; 15% perlite 2-6 mm). Each pot’s upper surface was covered at soil level by a round piece of plastic sheet (green or transparent according to the treatment, Fig. 1 and Supplementary Materials, Fig. S1). A coconut fibre disk was inserted between the plastic sheet and the soil to reduce warming and to improve air circulation and evaporation. The plants were randomly arranged in a grid pattern in a horticultural tunnel covered by a hail protection net, at a minimum distance of 46 cm from each other to diminish light competition between the plants, and all plants were irrigated ad libitum. For each cultivar and treatment, four plants were randomly selected and harvested at physiological stages R2 (full bloom), R6 (full seed stage, characterized by green pods containing fully developed green seeds), and R8 (full maturity, marked by leaf abscission and pods attaining their mature colour; Fehr and Caviness 1977). Plant roots were washed, and shoots and roots dried (40 °C, min 48 hours) and weighed. At full maturity, soybean pods were threshed and plant level seed yield determined. No plants of the cultivar ‘Capital’ were collected at R2 due to insufficient planting material resulting from poor germination (Supplementary Materials, Table S2). During the early growth period, leaf damage caused by thrips was observed. Therefore, leaf damage was scored visually 36 days after sowing (09.07.2024), using an ordinal scale ranging from 0 (no damage) to 3 (deformed leaves). Afterwards, all plants were treated with insect-repellent and no additional thrips or thrips damages were seen.

**Figure 1.**
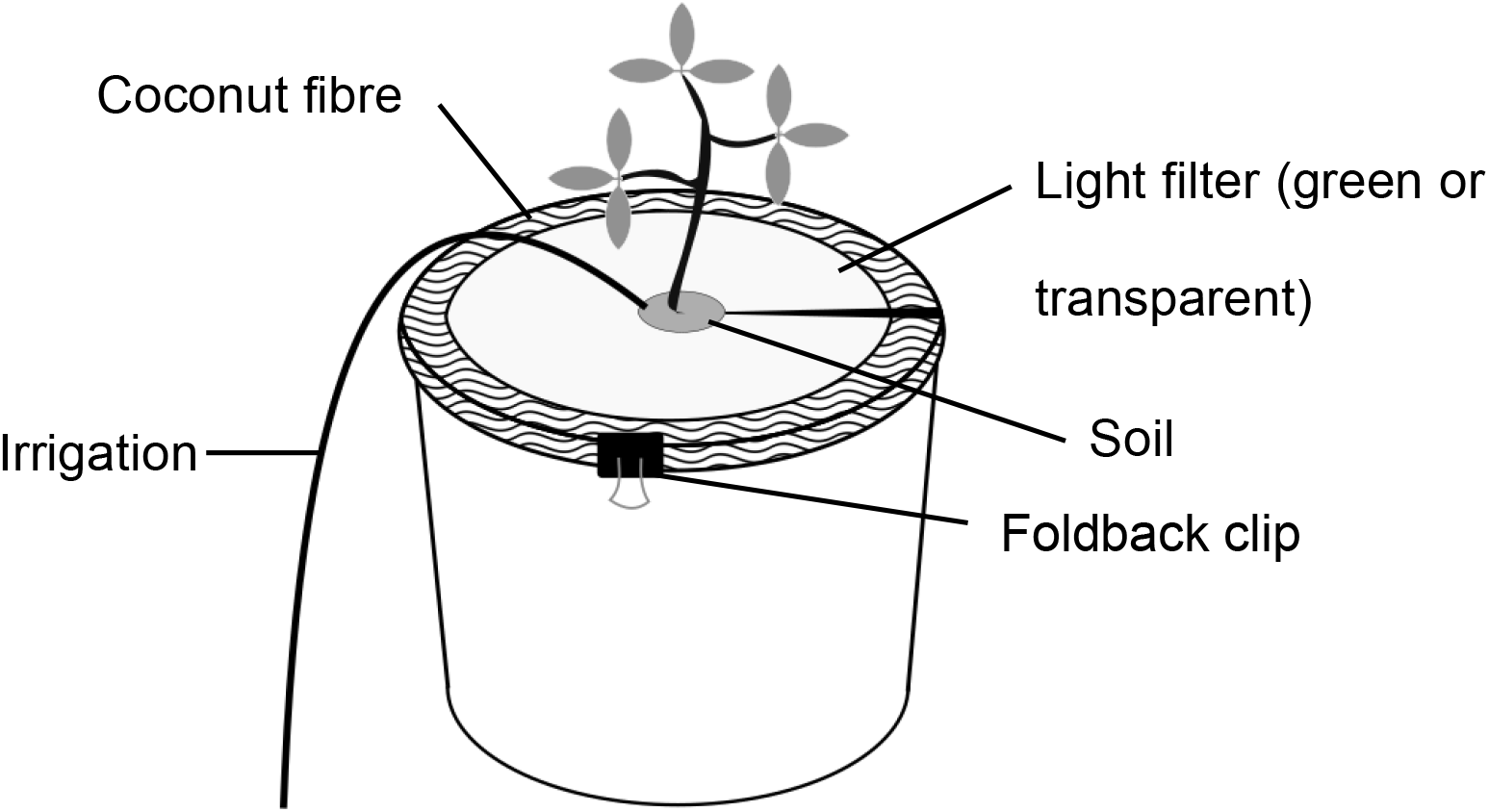
Schematic representation of the experimental setup. Each plant was cultivated in an individual pot, the soil surface of which was covered with a coconut fibre disc. A light filter was affixed directly to the upper surface of the disc. To ensure mechanical stability during field experiments, the disc–filter assembly was secured using foldback clips. The filter is either transparent, not altering the light wavelength composition; or green, reducing the amount of red and blue light reflected. Plants sense the presence of neighbours through shifts in light spectrum and might adapt their growth strategy accordingly (Supplementary Materials, Figure S1).

### Statistical analysis

All statistical analysis were performed in R version 4.4.3 (www.r-project.org). Multilevel linear models were fitted using the aov() function using its Error() option (version 4.5.0). Fixed effects included, in this order, year of release (YOR, continuous variable) and SAI treatment (categorical: shade-avoidance inducing or control), and the interaction of both terms. Error terms were cultivar, and the interaction of genotype with the SAI treatment. Effects of YOR therefore were tested using cultivar as error stratum, and the YOR x SAI treatment interaction used cultivar x SAI as error term. All dependent variables were natural log-transformed.

For the analysis of the herbivore damage, a cumulative-link mixed model was fitted using the *clmm2()* function from the *ordinal* package. Thrips damage was treated as an ordinal response, with the fixed and error terms as described above for the linear models.

## Results

### Plants under SAI treatment suffered from greater herbivore damage at early growth stage

After 36 days of growth (growth stage V4), we observed the presence of thrips in our experiment, often accompanied by early signs of leaf puncturing. Before treating against these unexpected pests, we visually quantified feeding signs on each plant individual. Plants growing under SAI treatment showed significantly more damage from thrips than plants under the control ratio treatment (Odds ratio = 0.356, P = 0.001; Fig. 2a), in line with known trade-offs between defence against herbivores and competition for light (Cipollini 2004; McGuire and Agrawal 2005). Also, modern cultivars were more susceptible to thrips than historical cultivars (Odds ratio = 1.006, P = 0.025; Fig. 2b). After treating plants against thrips, no damage was observed on new leaves until the end of the experiment.

**Figure 2.**
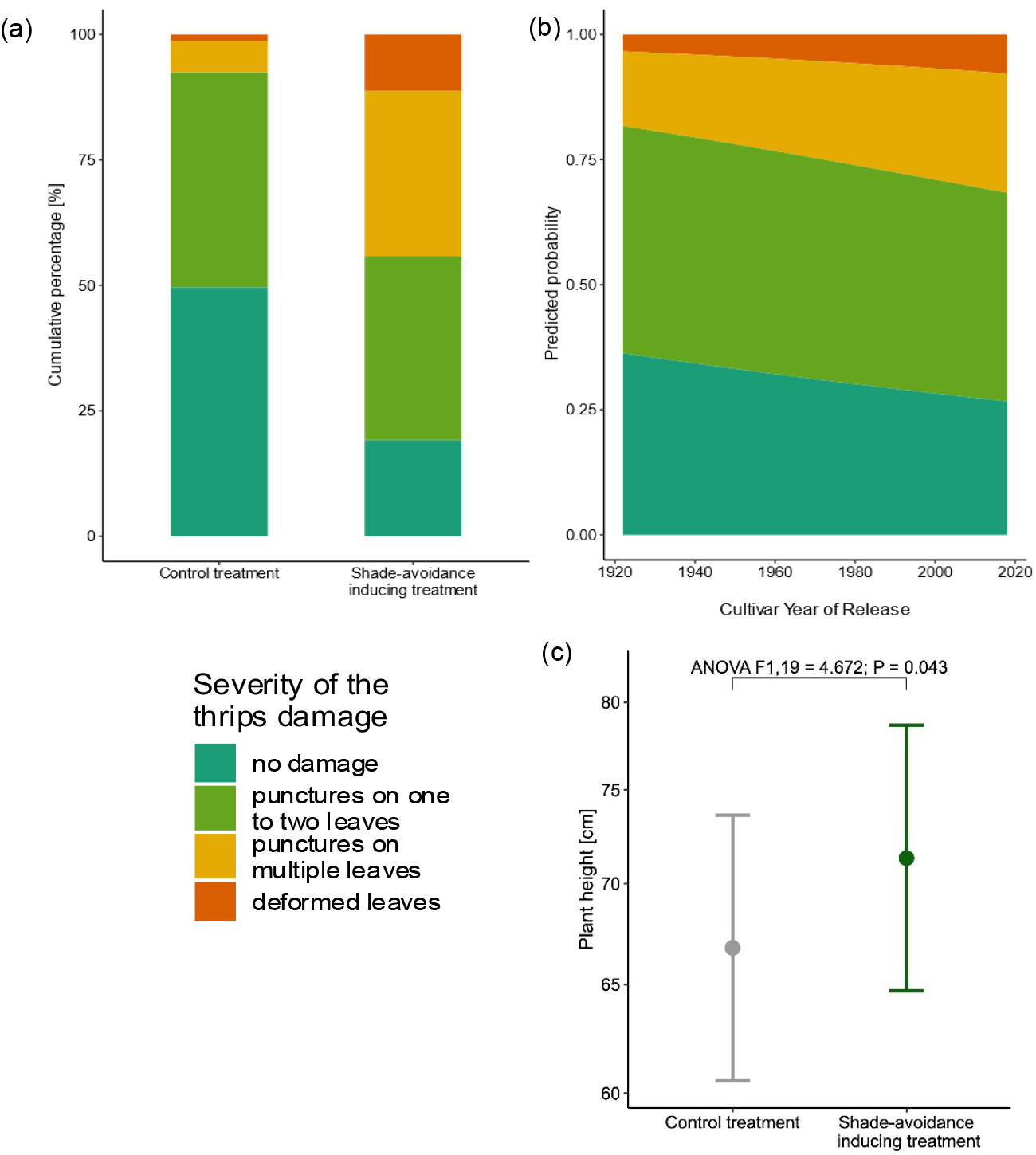
(a) Comparison of the thrips damage at growth stage V4 between the two treatments as cumulative percentage of the four severity categories. Plants under the shade-avoidance inducing treatment suffer higher thrips damages than control plants (b) Predicted probability of infestation over the cultivar year of release. More modern cultivars show higher thrips damages than historic cultivars. (c) Although the plants in the shade-avoidance inducing treatment at growth stage V4 showed higher susceptibility for thrips damages, at full height (growth stage R6) they were significantly taller compared to the control treatment, suggesting a shade-avoidance elongation response. The points and error bars represent the estimated marginal means from the mixed effects model ± SE. Plant height is depicted on the logarithmic scale.

At full height (growth stage R6), plants were taller in the SAI than in the control treatment (ANOVA F_1,19_ = 4.672, P = 0.043; Fig. 2c), in accordance with known shade avoidance responses. This suggests that the effects observed in the later growth stages are indeed due to shade avoidance responses and not to result of earlier leaf damage by thrips, since the latter would have reduced growth.

### Modern cultivars are more productive and differ in shoot architecture

Seed production significantly increased with year of release of the cultivars (ANOVA F_1,19_ = 6.594, P = 0.018), indicating breeding progress (Figure 3). At the same time, the number of branches per plant decreased with year of release (ANOVA F_1,19_ = 10.97, P = 0.003). Hence, modern cultivars did not produce more seeds by producing a greater number of equally productive branches, but by increasing the productivity per branch. Additionally, no change in stover production over time was observed, indicating that vegetative biomass (shoot biomass excluding yield) did not increase through breeding (Supplementary Materials, Table S3). However, modern cultivars tended to be more compact than historical cultivars, shown by an increasing ratio of shoot biomass to plant height over the last century (ANOVA F_1,19_ = 3.3, P = 0.085). This compactness may contribute to improved plant stability or resource use efficiency. Thus, breeding successfully led to an increase of plant productivity over time, while simultaneously changing the cultivars’ architecture.

**Figure 3.**
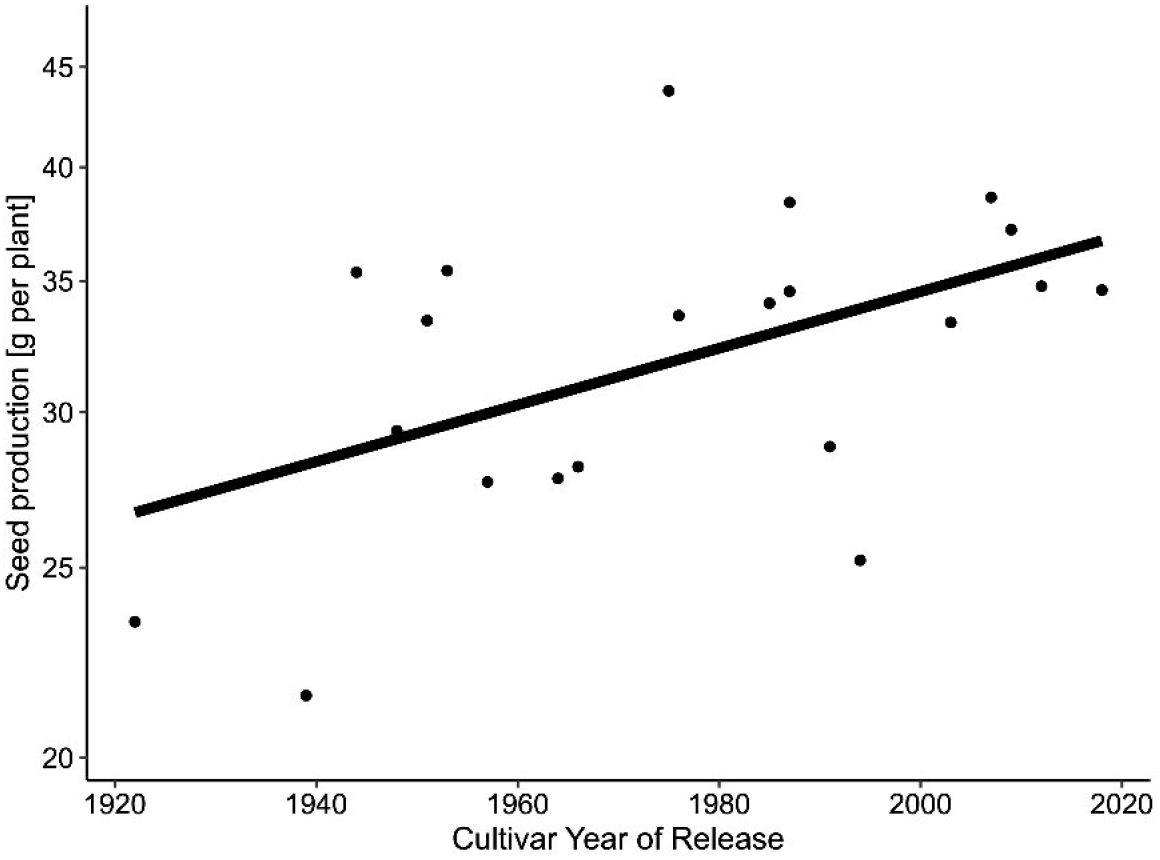
Cultivars released from 1922 to 2018 showed a significant increase in seed productivity at individual plant level (ANOVA F_1,19_ = 6.594, P = 0.018). The points correspond to the mean seed production (depicted on a logarithmic scale, *n* = 8 for each cultivar) of the 21 soybean cultivars used in the experiment.

Finally, breeding progress of seed production was higher in control than under the SAI treatment (interaction of YOR x treatment on seed production; ANOVA F_1,19_ = 4.771, P = 0.041; Fig. 4a). This progress was, however, not due to increased total biomass, but to a large degree due to an increased allocation of biomass to seeds. Indeed, the change of allocation over breeding history was lower under the SAI treatment (interaction of YOR x treatment on seed production - to - shoot biomass ratio; ANOVA F_1,19_ = 6.339, P = 0.021; Figure 4b).

**Figure 4.**
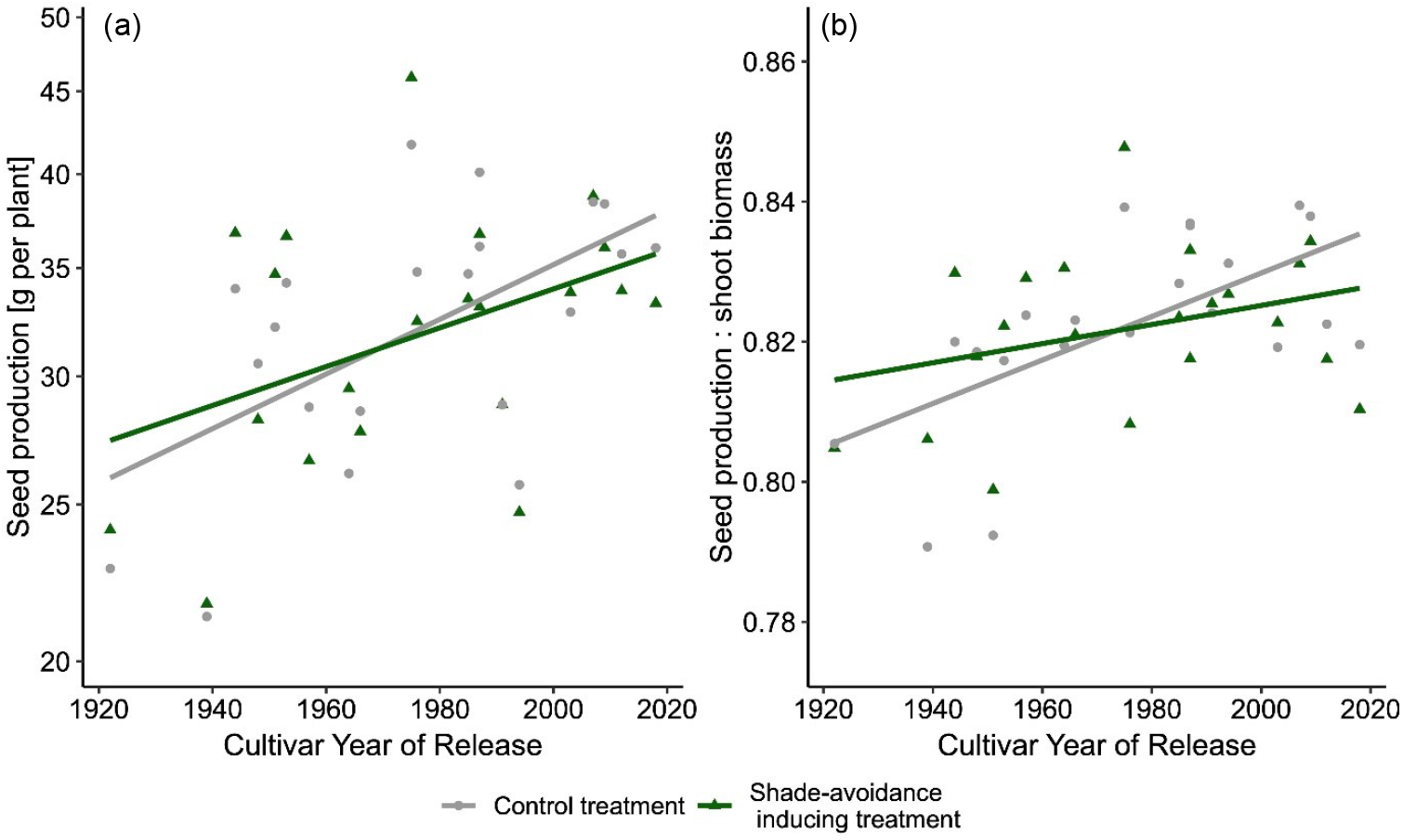
(a) The breeding progress was higher in the control treatment than in the shade-avoidance inducing treatment, as shown by the steeper slope of the grey line compared to the green one (ANOVA F_1,19_ = 4.771, P = 0.041). The grey dots correspond to the mean seed production of a cultivar under the control treatment (*n* = 4), whereas the green triangles are the mean seed production per cultivar under the SAI treatment (*n* = 4). Values depicted on a logarithmic scale (b) The control treatment shows a steeper increase towards a more favourable seed production to shoot biomass – ratio (ANOVA F1,19 = 6.339, P = 0.021).

## Discussion

We investigated the effects of altered light conditions on growth and biomass allocation in soybean cultivars released over nine decades of Canadian breeding history. We found plants exposed to simulated light competition to be more susceptible to herbivore damage than control plants, consistent with expectations due to shade avoidance responses (McGuire and Agrawal 2005; Ballaré and Pierik 2017). Moreover, more recently released cultivars were more susceptible to herbivores than older ones. At the same time, we found significant breeding progress for seed production, with modern cultivars producing more seeds than historical cultivars. However, the rate of this increase was steeper when light quality was not manipulated than when a SAI treatment mimicked the presence of neighbours. This increase was mostly driven by an improved allocation of resources to seeds over time, but with diminishing rate increases under shade avoidance treatments.

### Modern cultivars are both more productive and closer to the soybean ideotype than older cultivars

Here, we confirmed that breeding progress over decades led to higher seed production in soybean. Additionally, modern cultivars show morphological changes in agreement with a soybean ideotype, characterized by a reduced number of branches, short internode length, and erect leaves, as well as other characteristics that enhance light capture efficiency at high planting densities (Kokubun 1988; Lyu et al. 2023). However, we also observed higher susceptibility to herbivore damage in more modern cultivars – as well as plastic responses to simulated shading. The simultaneous increase in herbivore susceptibility and improvement in ideotype traits may reflect complex underlying mechanisms. While some shade avoidance responses may support canopy optimization (Zhou et al. 2024), their persistence, despite apparent group-level costs, might arise because they confer individual-level advantages during early breeding, where selection favours competitive success rather than collective performance. It is also worth noting that while ideotypes are generally defined based on fixed morphological traits, many relevant responses, such as response to shading, are at least partially plastic. This suggests that future breeding efforts may benefit from developing “plastic ideotypes” that show attenuated responses to environmental cues such as light quality changes, rather than focusing solely on static traits.

### Plants exposed to simulated light competition show increased herbivore susceptibility

Multiple studies have shown trade-offs between competition and herbivore susceptibility (McGuire and Agrawal 2005; Pellissier et al. 2018), a special case of the wider growth-defence trade-off (Herms and Mattson 1992; Fiorucci 2020). In our experiment, plants grown under SAI treatment, simulating the presence of neighbors through changes in light spectrum, exhibited increased thrips damage. While we did not quantify defence-related traits directly, this observation is consistent with previous findings showing that shade avoidance responses are accompanied by reduced investment in defence pathways (Cerrudo et al. 2012, 2017; Ballaré 2014). In natural ecosystems, rapid elongation to outgrow neighbours can increase light access and individual fitness, but under herbivore pressure, reduced defence investment may be costly. In crops, especially in monocultures, this effect may be disadvantageous for two reasons. First, if all plants react similarly to the competition cues and elongate to escape the canopy, this might result in direct yield losses due to a suboptimal resource allocation. Second, plants reacting to competition cues by reducing defence could suffer increased pest damage, leading to indirect yield losses not directly caused by competition. From a breeding perspective, it may be worth considering how responsiveness to SAI cues interacts not just with architecture and yield, but also with susceptibility to biotic stress. In this context, selecting for reduced sensitivity to SAI signals could have multiple benefits, including reduced pest susceptibility. However, unlike in cereal crops where shade avoidance has been reduced partly by selecting for shorter plants, a similar approach may not be viable in soybean. This is because soybean yield depends strongly on internode number, which is typically reduced in shorter plants (Liu et al. 2020). Experimentally, however, height and shade avoidance reductions have indeed been shown to be associated with relatively large yield benefits, especially under dense planting and at northern latitudes (Qin et al. 2023).

### Breeding improved soybean cultivars seed production, but additional potential might exist

Soybean yield has greatly increased in the last century, and the genetic gain through breeding progress has been documented in multiple studies (Luedders 1977; Specht et al. 1999; Cober et al. 2005; Kahlon et al. 2011). Here, the significant interaction between year of release and treatment on seed allocation suggests that breeding progress has been smaller under SAI conditions than in the control. Field studies comparing genetic gain in soybean under varying planting densities suggest that modern cultivars tolerate high stand densities, i.e. high within-stand competition, better than older cultivars (Cober et al. 2005; Suhre et al. 2014), but the evidence to support this claim is not very strong (De Bruin and

Pedersen 2009). Two possible explanations may account for these findings. First, it is possible that the plastic responses we observe under SAI treatment have a beneficial impact on population level yield at high densities, for example by optimizing canopy orientation or plants’ architecture. If so, these traits may have been retained during selection and may not limit yield at the group level. Alternatively, the stronger increase under control conditions may reflect that early-stage breeding exposed plants to competitive environments, but selected individuals based on their own performance. This likely favoured traits such as shade avoidance, which improve individual competitiveness but may be detrimental at the group level under dense planting (Keuskamp et al. 2010; Bongers et al. 2018; Chen et al. 2023). This would imply that current cultivars may still perform suboptimal in dense stands because traits like shade avoidance were selected for individual success in early competitive breeding stages, without regard for group-level performance. Targeted selection for group performance under high-density conditions, focused on population yield rather than individual competitiveness, could therefore reduce maladaptive shade responses and further improve yield.

## Conclusion

While traditional breeding has successfully improved soybean yield and ideotype traits, this study highlights that persistent shade avoidance responses, possibly favoured due to increased individual competitiveness in early breeding stages, can constrain performance and increase herbivore susceptibility when light wavelength composition is altered, as is typical in dense stands. To fully unlock soybean’s yield potential, future breeding efforts should focus on selecting for traits that enhance group-level performance under high-density conditions. This includes reducing potentially disadvantageous plastic responses to light competition and developing “plastic ideotypes” that integrate adaptive environmental responsiveness. Such strategies could lead to crops better optimized for modern agricultural systems, improving both productivity and resilience.

## Acknowledgements

We thank the USDA germplasm collection and Claude-Alain Bétrix (Agroscope) for providing seeds; Jürgen Krauss (Agroscope) for technical support. This study received funding from the Swiss National Science Foundation (grant no. 310030_192537 to SEW).

## Data availability

The dataset used in this study is available from the Zenodo data repository (DOI: 10.5281/zenodo.17183257)

## Supplementary material

**Figure S1.**
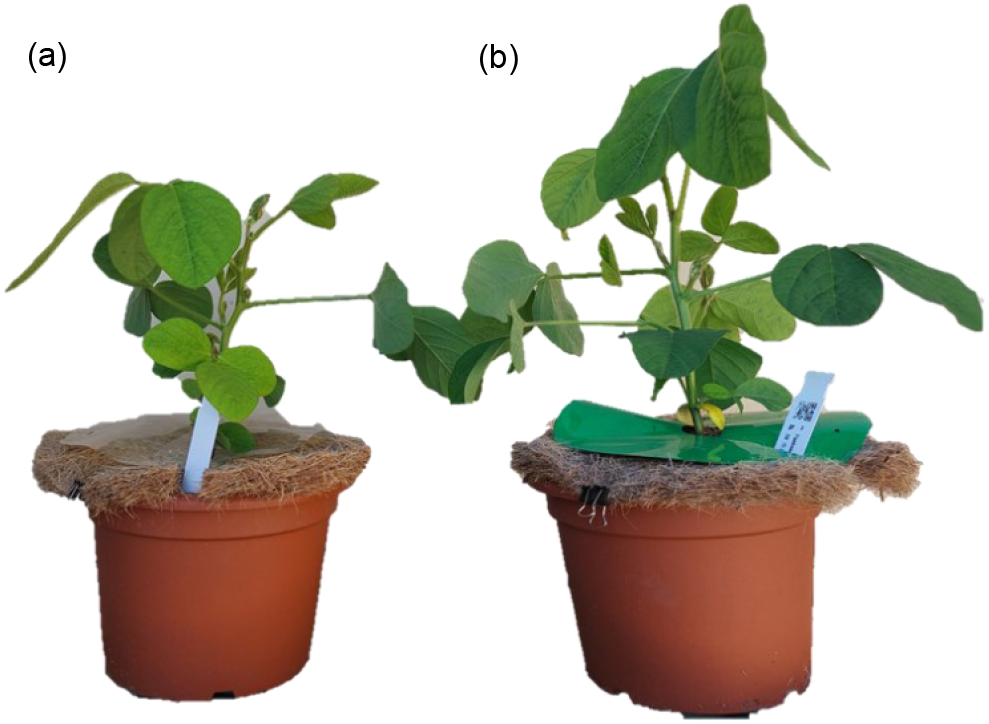
Experimental setup of the two treatments. Plant (a) is growing under the control treatment with a transparent filter above the coconut fibre disc. Plant (b) is growing under the SAI treatment with a green filter fixed on the coconut fibre disc.

**Table S1.**
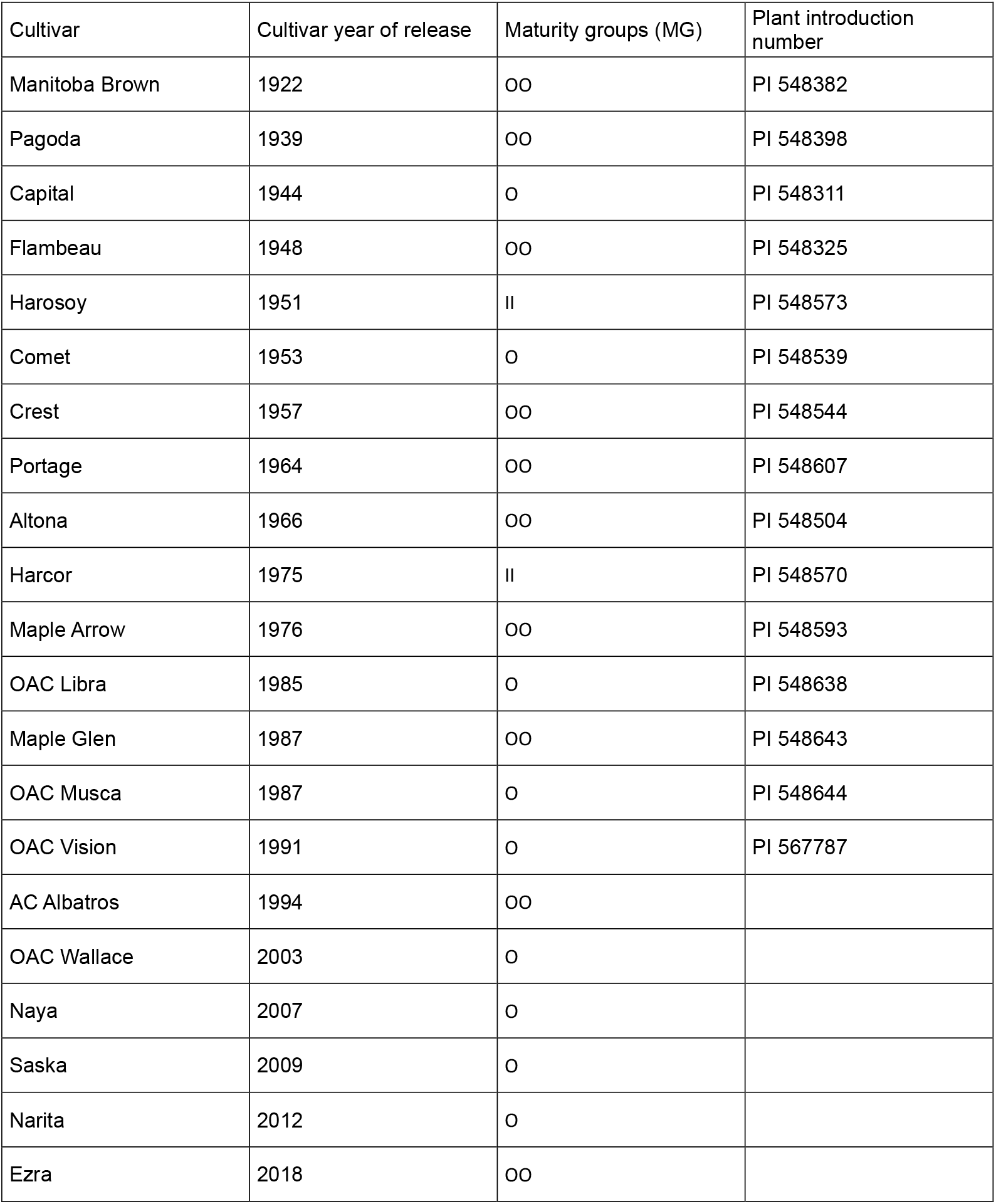
Description of cultivars utilized in this study. We used 21 early maturing cultivars from the Canadian breeding program, spanning almost a century of breeding.

**Table S2.** Harvest dates for the different growing stages and cultivars. We harvested all cultivars at the same phenological states, explaining the difference in harvest dates.

| Cultivar | Harvest VC | Harvest R2 | Harvest R6 | Harvest R8 |
| --- | --- | --- | --- | --- |
| Manitoba Brown | 12.06.2024 | 21.07.2024 | 15.08.2024 | 11.09.2024 |
| Pagoda | 12.06.2024 | 21.07.2024 | 28.08.2024 | 11.09.2024 |
| Capital | 12.06.2024 |  | 02.09.2024 | 11.09.2024 |
| Flambeau | 12.06.2024 | 21.07.2024 | 28.08.2024 | 29.09.2024 |
| Harosoy | 12.06.2024 | 21.07.2024 | 02.09.2024 | 29.09.2024 |
| Comet | 12.06.2024 | 21.07.2024 | 02.09.2024 | 29.09.2024 |
| Crest | 12.06.2024 | 21.07.2024 | 02.09.2024 | 11.09.2024 |
| Portage | 12.06.2024 | 21.07.2024 | 28.08.2024 | 11.09.2024 |
| Altona | 12.06.2024 | 21.07.2024 | 28.08.2024 | 11.09.2024 |
| Harcor | 12.06.2024 | 21.07.2024 | 02.09.2024 | 29.09.2024 |
| Maple Arrow | 12.06.2024 | 21.07.2024 | 02.09.2024 | 29.09.2024 |
| OAC Libra | 12.06.2024 | 21.07.2024 | 02.09.2024 | 29.09.2024 |
| Maple Glen | 12.06.2024 | 21.07.2024 | 02.09.2024 | 29.09.2024 |
| OAC Musca | 12.06.2024 | 21.07.2024 | 02.09.2024 | 29.09.2024 |
| OAC Vision | 12.06.2024 | 21.07.2024 | 28.08.2024 | 11.09.2024 |
| AC Albatros | 12.06.2024 | 21.07.2024 | 02.09.2024 | 11.09.2024 |
| OAC Wallace | 12.06.2024 | 21.07.2024 | 02.09.2024 | 11.09.2024 |
| Naya | 12.06.2024 | 21.07.2024 | 02.09.2024 | 29.09.2024 |
| Saska | 12.06.2024 | 21.07.2024 | 02.09.2024 | 29.09.2024 |
| Narita | 12.06.2024 | 21.07.2024 | 02.09.2024 | 29.09.2024 |
| Ezra | 12.06.2024 | 21.07.2024 | 02.09.2024 | 29.09.2024 |

**Table S3:**
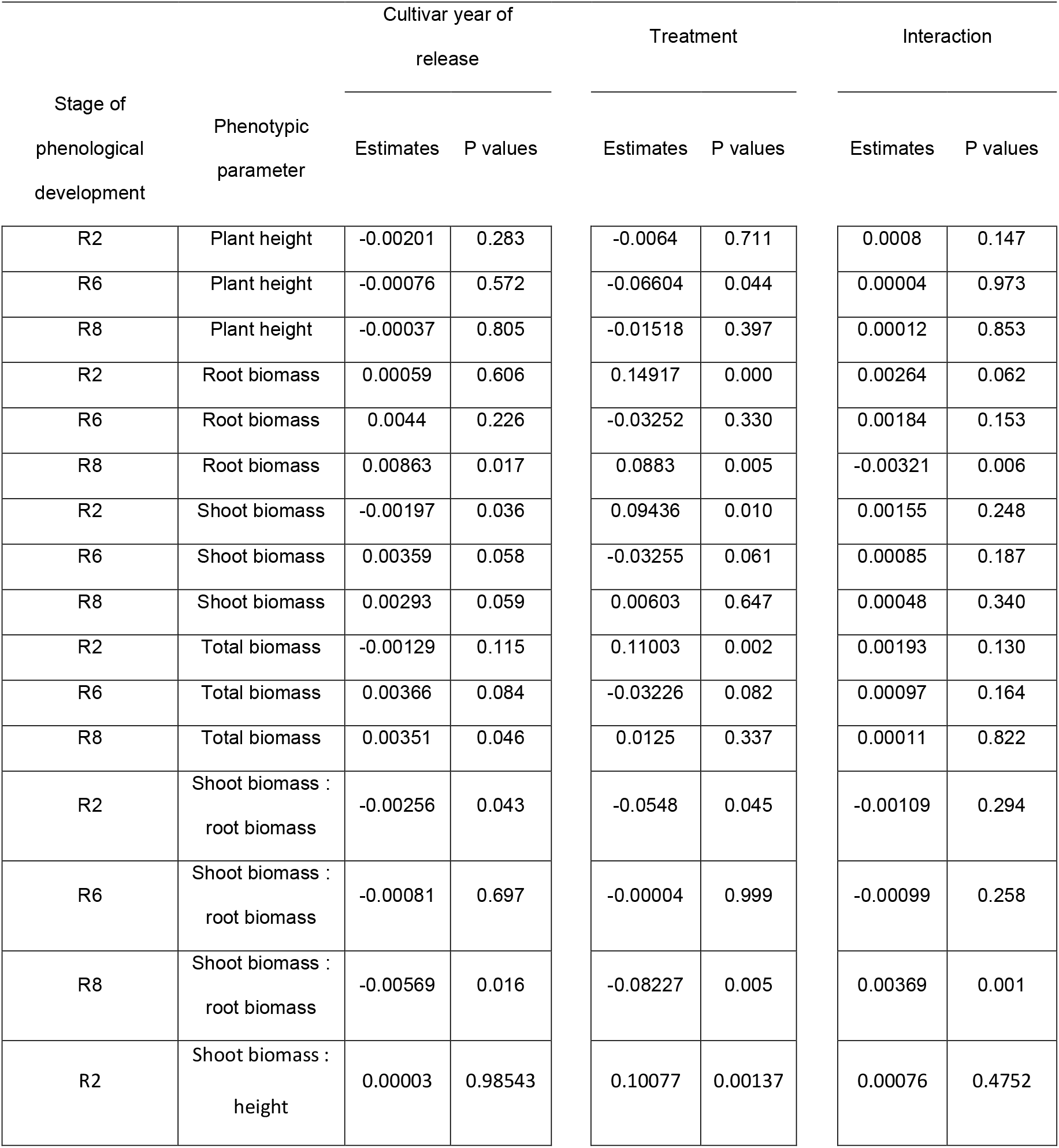
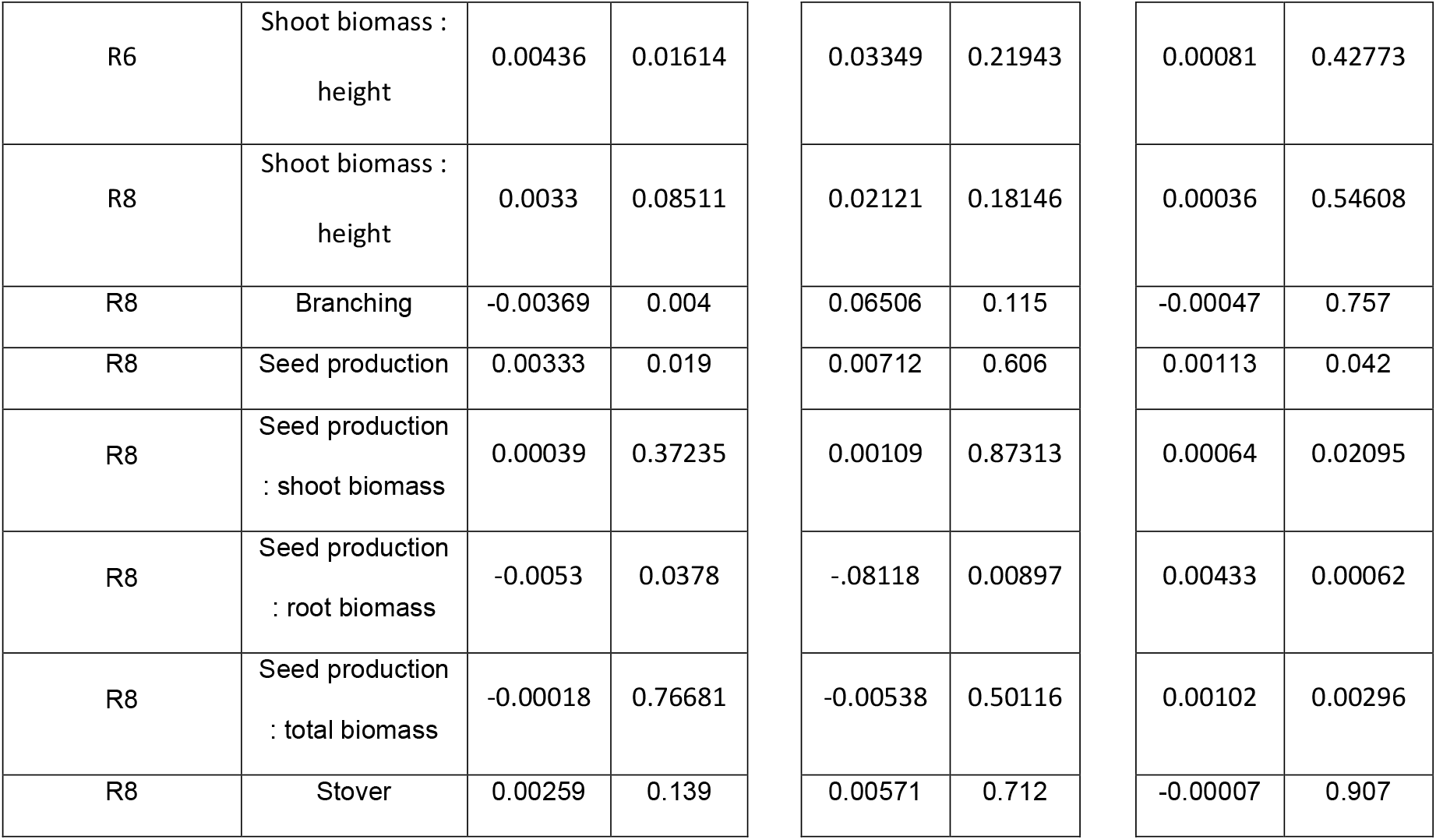
Estimates and p-values resulting from the linear mixed models. Shoot biomass is the aboveground biomass including shoots, leaves, pods, seeds etc. The total biomass is the sum of shoot biomass and root biomass. Stover is the shoot biomass minus the seed mass.

